# Non-cell autonomous small RNA silencing in female gametes and early embryo of Arabidopsis

**DOI:** 10.1101/2022.03.21.485150

**Authors:** JA Schröder, DMV Bonnet, PE Jullien

## Abstract

In recent years, small RNA movement has been both hypothesized and shown to be an integral part of the epigenetic DNA methylation reprogramming occurring during plant reproduction. It was suggested that the release of epigenetic silencing in accessory cell types or tissues is necessary to reinforce epigenetic silencing in the gametes (egg cell and sperm cells) which would in turn ensure the genomic stability of the next generation plant. Small RNA movement was indeed shown to occur during male gametogenesis. However, the situation within the female gametophyte and in early seed development is mostly unknown. Here, we show that small RNA can induce non-cell autonomous silencing from the central cell towards the egg cell but also from the synergids to the egg cell and central cell. In addition, we also observe a non-cell-autonomous silencing from the central cell or endosperm towards the early embryo. Our data shows, that in addition to the movement of sRNAs during pollen development, sRNA movement also occurs in the female gametes.

## Main text

In plants, small RNAs (sRNAs) can move long distances via the plant vasculature as well as cell-to-cell via plasmodesmata^1,2^. sRNA movement has been implicated in several processes, such as plant defence and plant development. Mobile sRNAs can regulate gene expression remotely either at the post-transcriptional or at the transcriptional level. Transcriptional silencing induced by sRNAs relies on a pathway called RNA-dependent DNA methylation (RdDM) which results in the *denovo* DNA methylation of the sRNAs complementary DNA sequence. Complex changes of the plant methylome occur during sexual reproduction likely to ensure the silencing of transposable elements in the embryonic lineage. During pollen development, mobile sRNAs were shown to be involved in this process. sRNAs move from the vegetative cell to sperm cells^3^ at pollen maturity and from the sporophytic tapetum to microspores^4^. In contrast to male gametogenesis, available data for sRNA movement in mature female gametes are limited to two reports. In the first, an artificial microRNA expressed in the central cell (CC) was shown to silence GFP expressed in the egg cell (EC)^5^. In the second, labelled sRNAs injected in the CC were shown to freely diffuse to the EC^6^. These two reports focused on sRNA movement specifically from the CC to the EC and other sRNAs movements were not investigated.

To assess additional types of sRNAs movement from the accessory cells of the female gametophyte towards adjacent cell types, sRNAs generating hairpins expressed under a CC promoter (*pDD65*) or under a synergid cells (Syn) promoter (*pDD35*) were transformed into a marker line expressing GFP ubiquitously (*LIG1:GFP*). As published, we confirmed that both promoters are specifically expressed in their respective cell types (CC for pDD65 and Syn for pDD35) and that their expression starts after cellularisation of the female gametophyte (Figure S1A-F). Additionally, to determine whether the enzymes necessary for the processing of hairpin precursors into sRNAs are present in the different cell types, we analysed the expression of the four *Arabidopsis* DICER (DCL) proteins using full locus reporter constructs. We could observe that all DCLs are expressed in the CC, although DCL1-mCherry and DCL4-GFP are expressed at a lower level than DCL2-GFP and DCL3-GFP (Fig S2J-M). A very high expression of DCL1-mCherry, DCL2-GFP, and DCL3-GFP was observed in the Syn. We conclude that DCLs are present in the mature female gametophyte to process hairpin precursors into sRNAs.

To analyse a potential non-cell autonomous silencing within the mature female gametophyte, we measured the mean fluorescence intensity of nuclei from the Syn, EC, CC, and integuments (Int) of transgenic lines homozygous for both the cell-specific hairpin construct (*pDD35hp* or *pDD65hp*) and *LIG1:GFP* at one day post emasculation (1DAE) (Figure 1A-D, H, I). In *pDD35hp* and *pDD65hp* lines, we observe GFP silencing in the Syn (Fig1C,H) and the CC respectively (Fig1D,I) showing the functionality of a cell-autonomous silencing. Interestingly, non-cell autonomous silencing was observed from the Syn towards both the EC and the CC in the *pDD35hp* lines (Fig1 C,H, FigS2A), strongly suggesting that sRNAs can move from the Syn toward the EC and CC. In neither of two independent experiments, could we detect a movement from the Syn to the Int (Fig1 H, FigS2 A). As published, we saw a non-cell autonomous silencing in the EC in our *pDD65hp* lines (Fig 1D,I and FigS2B). However, intriguingly, silencing movement was neither observed between the CC and the Syn nor towards the Int. The absence of silencing movement between the CC and Syn could result from either the existence of a transport checkpoint between these two cell types or from a difference in the quantity of sRNAs generated by each construct. Indeed, *pDD35* seems to be expressed at a higher level compared to *pDD65* (Fig S1 G,F). The higher expression of *pDD35hp* together with the strong expression of DCLs in the Syn could therefore generate more sRNAs with mobile silencing potential than are produced in the *pDD65hp* lines. The cell-to-cell movement of sRNAs occurs mostly by symplastic connections called plasmodesmata. Despite controversial literature^7,8^, plasmodesmata connections have been reported between all the cells of the mature female gametophyte, suggesting that the mature female gametophyte represents one common symplast. We therefore conclude that beyond a CC to EC movement, sRNAs move freely within the mature female gametophyte prior to fertilization.

**Figure. 1.**
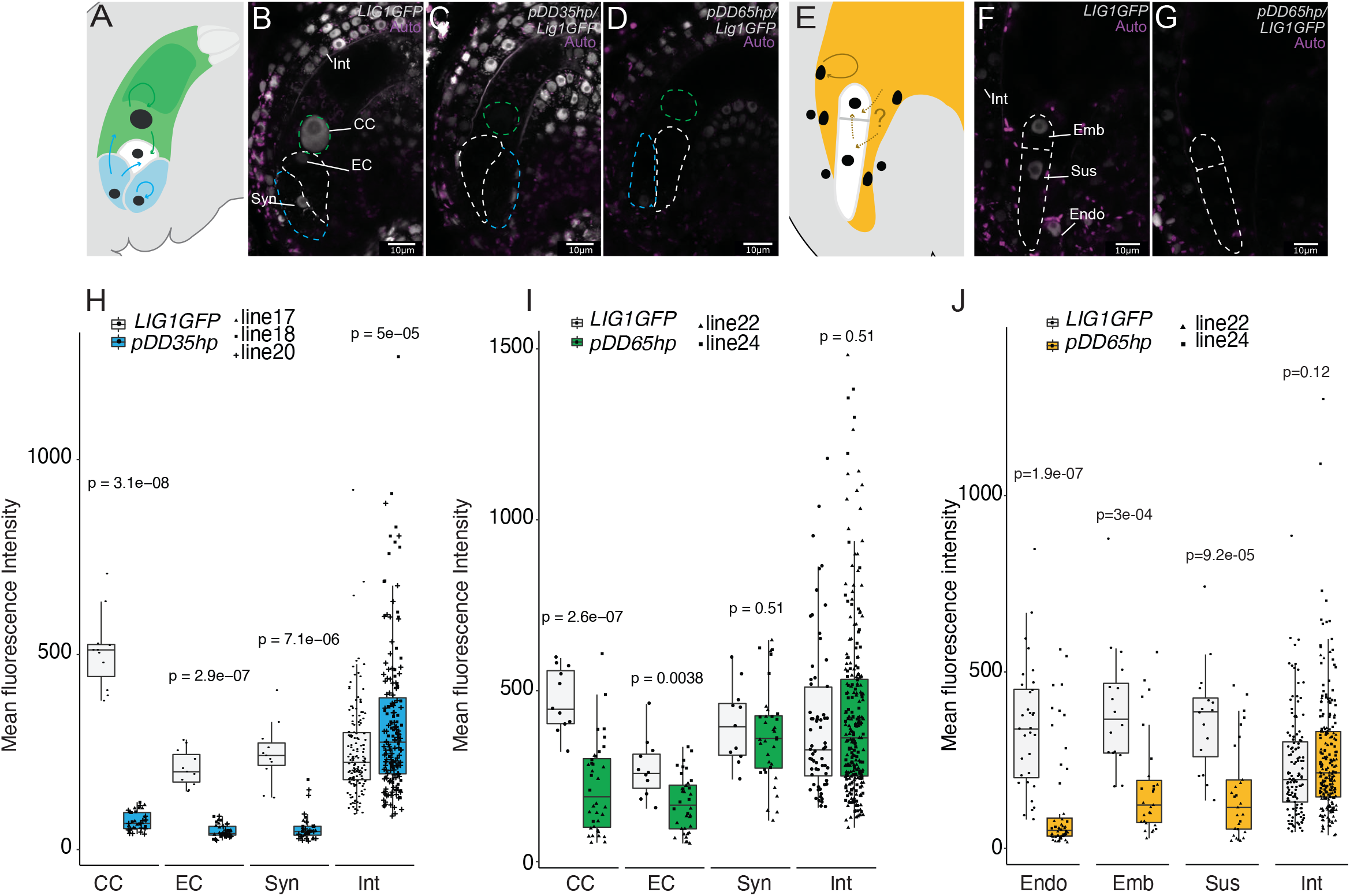
small RNA movement within the female gametophyte and early seed. (A) Schematic representation of an Arabidopsis ovule and the intercellular movement observed in this study. The central cell where the *pDD65* promoter is active is coloured in green while the synergides where the *pDD35* promoter is active are coloured in blue. (B-D) Confocal images of ovules showing autofluorescence (Auto) in magenta and LIG1:GFP expression in grey scale of *LIG1GFP* control (B), *pDD35hp;LIG1GFP* (C) and *pDD65hp;LIG1GFP* (D) ovules. Dashed lines highlight the egg cell (EC), synergide cells (Syn), and the central cell nucleus (CC). (E) Schematic representation of an *Arabidopsis* seed and the intercellular movement observed in this study. The endosperm where the pDD65 promoter is active is coloured in orange (F-G) Confocal images of seeds showing the autofluorescence (Auto) in magenta and LIG1:GFP expression in grey scale of *LIG1GFP* control (F), and *pDD65hp;LIG1GFP* (G) seeds. Dashed lines highlight the 1 day after fertilization embryo proper (Emb), suspensor (Sus), endosperm (Endo) and the Integument (Int) nuclei. (H-J) Box plots showing the quantification of the mean fluorescence intensity measured before fertilisation (one day after emasculation, 1DAE) for *pDD35hp;LIG1GFP* (H) and for *pDD65hp;LIG1GFP* (I) as well as after fertilisation (one day after fertilisation, 1DAP) for *pDD65hp;LIG1GFP* (J). *LIG1GFP* plants were grown in parallel and quantified as a control. Different symbols represent independent transgenic lines for *pDD35hp;LIG1GFP* (abbreviated *pDD35hp*) and *pDD65hp;LIG1GFP* (abbreviated *pDD65hp*). Controls of the *pDD35* and *pDD65* promoter activity can be found in FigS1A-I. Additional measurement from additional time point and independent experiments can be found in FigS2. The scale bar indicates 10µm.

A hypothesis that is often mentioned in papers and reviews is the potential movement of sRNA from the fertilized endosperm to the embryo to direct the increase in DNA methylation observed during embryo development. Such movement, if occurring, would have to happen at early stages of embryo development before the embryo becomes clearly symplastically and apoplastically isolated from the endosperm^9^. To test whether we could observe a silencing movement from the endosperm to the embryo, we quantified the mean fluorescence intensity in the different cell types of the developing seed (endosperm, embryo, suspensor and integuments) at different time-points of early seed development in the *pDD65hp;LIG1GFP* lines (Fig1E-G, J and FigS2). We first confirmed that *pDD65* is not active in the embryo but solely in the endosperm (Fig S1G-I) and showed that all DCLs are expressed in the endosperm except *DCL4-GFP* (Fig S1N-Q). GFP fluorescence intensity was clearly reduced in both the endosperm (cell-autonomous silencing) and the embryo/suspensor (non-cell-autonomous silencing) at one day after pollination (1DAP) (Fig1E-G, J). As expected, no silencing was observed in the integuments due to the absence of symplastic and apoplastic connections between the integument and the inside of the seed. GFP silencing in the embryo was less pronounced at 2DAP than at 1DAP and was not longer significant at 3DAP (Fig S2C). Our results show that a non-cell autonomous silencing occurs between the endosperm lineage and the early embryo but is rapidly lost during seed development. Nonetheless, using dyes, a loss of symplastic transport has been proposed to occur very shortly after fertilization^6,10^. Non-autonomous embryo silencing that we observe could also result from an inheritance of the central cell to egg cell silencing that we observed at 1DAE. A deeper analysis of the presence of plasmodesmata in the early zygote or in the basal cell of the suspensor as performed in other plants^11^ should help to distinguish between those two possibilities.

## Acknowledgements

We would like to thank Jasmin Sekulovski for her support concerning plant growth. We thank Gwyneth Ingram, Daniel Bouyer and Stefan Grob for critical reading of the manuscript. We acknowledge the support of the Microscopy Imaging Centre (MIC) from the University of Bern. PEJ, DMVB and JAS are supported by an SNF professorship grant (no.163946) attributed to PEJ.

## Author contributions

PEJ designed the study. JAS and DMVB performed the experiments. JAS conducted the main analyses. PEJ wrote the manuscript with support from JAS and DMVB.

## Supplemental information

### Material and Methods

#### Plant material

Arabidopsis thaliana Col-0 seeds were provided by the Nottingham Arabidopsis Stock Center (NASC). and were used as a wild-type background for all experiments. The *pLIG1:LIG1-GFP* line (abbreviated *LIG1GFP*) was previously described by Andreuzza *et al*.^1^. The DICER translational reporters lines: *pDCL1:DCL1-mCherry;dcl1 pDCL2:DCL2-GFP;dcl2-1, pDCL3:DCL3-GFP* and *pDCL4:DCL4-GFP;dcl4-2* were previously described by Pumplin N. *et al*. ^2^. Seeds were stratified at 4°C for two days and subsequently germinated and grown under long-day conditions (16h light 22°C / 8h dark 19°C) in a controlled growth chamber.

#### Cloning and transformation

Multisite Gateway Cloning technology (ThermoFisher Scientific) was used to obtain all constructs used during this study. Briefly, the promoters-CDS-Hairpin sequences were amplified using the Phusion High-Fidelity DNA Polymerase (Thermo) using the following primers : PJ-0634-AttB4-DD65 (GGGGA CAACT TTGTA TAGAA AAGTT GCTAA TCAAA ATTTA ACATT TAAAT AAA), PJ-0635-AttB1r-DD65 (GGGGA CTGCT TTTTT GTACA AACTT GCATCC TTTTC TACTTT GTTTTT GTTT), PJ-0622-AttB4-DD35 (GGGGA CAACT TTGTA TAGAA AAGTT GCTTC TGATA AAATG GAAAA TTCAA AGA), PJ-0623-AttB1r-DD35 (GGGGA CTGCT TTTTT GTACA AACTT GCGAG AAACA ATGGT GGCCA TTTTG TT), PJ-0022-AttB1-GFhp (GGGGA CAAGT TTGTA CAAAA AAGCA GGCTT AATGA GTAAA GGAGA AGAAC TTTTC A), PJ-0025-AttB2-GFhp (GGGGA CCACT TTGTA CAAGA AAGCT GGGTA CGTCC TCCTT GAAAT CGATT CCCTT), PJ-0023-AttB1-FPhp (GGGGA CAAGT TTGTA CAAAA AAGCA GGCTT ACAAG TTTGA GGGAG ACACC CTCGT C), PJ-0026-AttB2-FPhp (GGGGA CCACT TTGTA CAAGA AAGCT GGGTA TTAGTG GTGGTG GTGGTG GTGTT TG), PJ-0021-AttB1-H2B (GGGGA CAAGTT TGTACA AAAAAG CAGGCT TAATGG CGAAG GCAGA TAAGA AACCA G), PJ-0024-AttB2-H2B (GGGGA CCACT TTGTA CAAGA AAGCT GGGTA AGAAC TCGTA AACTT CGTAA CCGCC), PJ-0236-AttB1-DCL3 (GGGGA CAAGT TTGTA CAAAAA AGCAGGC TTAATG CATTCG TCGTTG GAG), PJ-0237-AttB2-DCL3 (GGGGAC CACTT TGTAC AAGAA AGCTG GGTAC TACTT TTGTA TTATG AC), PJ-0238-AttB4-pDCL3 (GGGGA CAACT TTGTA TAGAA AAGTTG CTGATC TTTGGG AGACGT AA), PJ-0239-AttB1r-pDCL3 (GGGGA CTGCT TTTTT GTACAA ACTTGC GACGAAC GGATA AAAAG G). The obtained PCR products were cleaned up using the GENEjET Gel Extraction or DNA Cleanup Micro Kit (Thermo) and subsequently recombined into a Donor vector using BP Clonase (Thermo). The obtained Entry vectors were subsequently recombined with the Destination vectors to obtain the final Expression vectors. The *pDD35:H2B-Clover* and *pDD65:H2B-Clover* constructs were recombined using pB7m34GW as destination vector^3^. The *pDD35:GFhp* and *pDD65:GFhp* were assembled using pBm42GWIWG8,1 as destination vector (https://gatewayvectors.vib.be). All plasmids were verified using colony PCR, digestion, and sequencing. Stable transgenic lines were obtained using the floral dipping method^4^ in *LIG1GFP* background. For each construct, a minimum of 8 independent insertion lines were observed to ensure consistent results. Confocal imaging and quantification were performed on at least two independent stable lines per experiments that were identified to be single insertion homozygous line according to the antibiotic resistance apart from *pDD35hp;LIG1GFP* line number 17 which was double insertion.

#### Microscopy

All microscopy experiments were performed using a laser scanning microscope (Leica SP5). For ovule experiments, plants were emasculated one day prior observation (1 day after emasculation, 1DAE). For seed observations, flowers were emasculated and one day later hand pollinated. Seeds were observed between one to three days after pollination (1-3 DAP). GFP images for all silencing experiments of Figure 1 and S2 were taken with the same confocal settings: 20% Argon Laser power, 288nm excitation, 498-565 nm for emission, gain of 100 and pinhole set to 1. For expression patterns of Figure S1, the laser power, gain, and emission bandpass were adapted depending on the stage and signal intensity of the different lines. If necessary for illustration purposes, brightness and/or contrast was uniformly modified using the Fiji installation of ImageJ^5^.

#### Image quantification

Quantification of the mean GFP fluorescence intensity was performed by using Fiji installation of ImageJ^5^. Due to the nuclear localization of the GFP signal of LIG1GFP, quantification of the mean GFP fluorescence intensity was performed on nuclei. Nuclei were selected manually based on the brightfield image and/or enhanced GFP or Autofluorescence channels. For mature female gametophyte quantification, the mean fluorescence of one nucleus per cell type was measured on the same picture and same confocal plane. For 1 DAP seeds, one embryo, one suspensor and two endosperm nuclei were quantified on the same picture and same confocal plane. At a later stage, due to the increased seed thickness, two pictures were taken of each seed: one for the endosperm and one for the embryo and suspensor. From 2 DAP seeds, two embryo nuclei, one suspensor nuclei and two endosperm nuclei were measured per analyzed seed. Due to a high variability in LIG1GFP fluorescence intensity in the integument (likely due to cell-cycle control of the DNA ligase 1), Inner-integument measurements were done on 7 nuclei situated either next to the central cell/synergids or next to the embryo/suspensor. A minimum of nine ovules or seeds from a minimum of two independent pistils or siliques and two to three independent transgenic lines were used.

## Supplementary Figure legends

**Figure S1.**
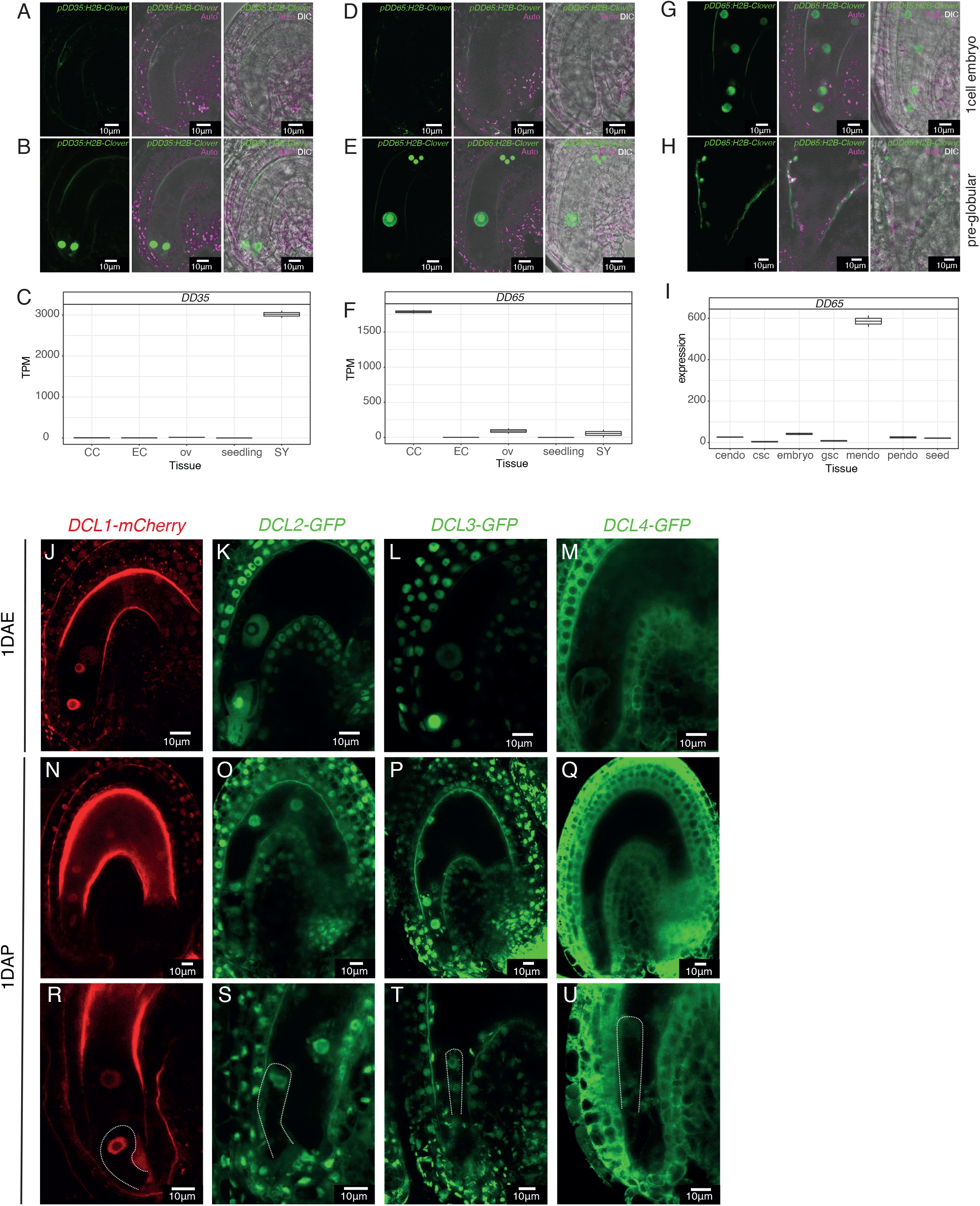
Promotor activity of pDD35 and pDD65 and DICER translational fusion expression pattern. (A-B) *DD35* promoter activity in mature synergids is illustrated by confocal images taken during gametogenesis stages FG5 (A) and FG6 (B) according to Christensen *et al*.^6^ using *pDD35:H2B-Clover* lines. The pictures represent: on the left the Clover signal (green), in the middle the merge of Clover with the autofluorescence signal (“Auto” in magenta), on the right the merge of Clover, Auto and the DIC (Differential interference contrast). (C) *DD35* RNA expression level plotted using publicly available single-cell transcriptomic data of mature female gametophytes^7^. (D-E) *DD65* promoter activity in the central cell is illustrated by confocal images taken during gametogenesis stages FG5 (D) and FG6 (E) using *pDD65:H2B-Clover* lines. The pictures represent: on the left the Clover signal (green), in the middle the merge of Clover with the autofluorescence signal (“Auto” in magenta), on the right the merge of Clover, Auto and the DIC (Differential interference contrast). (F) *DD65* RNA expression level plotted using publicly available single-cell transcriptomic data of mature female gametophytes^7^. (G-H) *DD65* promoter activity in the endosperm is illustrated by confocal images taken at 1DAP (G) and 2DAP (H) using *pDD65:H2B-Clover* lines. The pictures represent: on the left the Clover signal (green), in the middle the merge of Clover with the autofluorescence signal (“Auto” in magenta), on the right the merge of Clover, Auto and the DIC (Differential interference contrast). The embryo outline can be seen on the right image. (I) *DD65* RNA expression level plotted using publicly available pre-globular stage seed LCM microarray data^8^. (J-M) Confocal pictures of mature female gametophyte at 1DAE displaying the expression pattern of *pDCL1:DLC1-mCherry* (J), *pDCL2:DLC2-GFP(K), pDCL3:DLC3-GFP(L) and pDCL4:DLC4-GFP* (M). (N-Q) Confocal pictures of early seeds at 1DAP displaying the expression pattern of *pDCL1:DLC1-mCherry* (N), *pDCL2:DLC2-GFP(O), pDCL3:DLC3-GFP(P) and pDCL4:DLC4-GFP* (Q). (R-U) Confocal pictures of early seeds at 1DAP displaying the expression of DCLs in the early embryo of *pDCL1:DLC1-mCherry* (R), *pDCL2:DLC2-GFP(S), pDCL3:DLC3-GFP(T) and pDCL4:DLC4-GFP* (U). The scale bar indicates 10µm. The following abbreviations are used: central cell (CC), egg cell (EC), synergids (SYN), ovule (ov), chalazal endosperm (cendo), chalazal seed coat (csc), general seed coat (gsc), micropylar endosperm (mendo), peripheral endosperm (pendo) and Transcripts per million (TPM).

**Figure S2.**
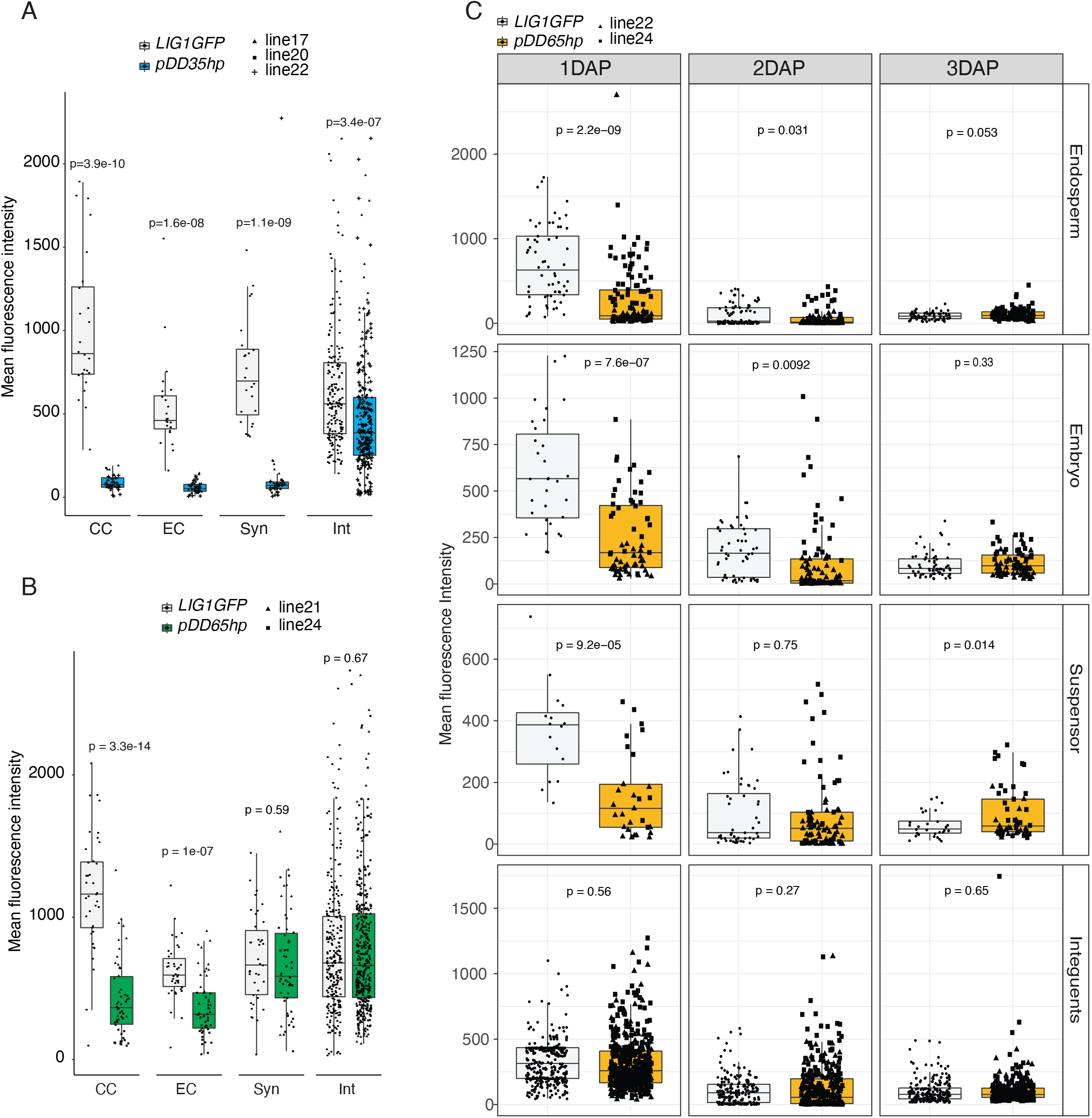
(A-B) Box plots showing the quantification of the mean fluorescence intensity measured before fertilisation (one day after emasculation, 1DAE) for *pDD35hp;LIG1GFP* (A) and for *pDD65hp;LIG1GFP* (B) from an independent experiment of the one in Fig1H-I. (C) Box plots showing the quantification of the mean fluorescence intensity measured after pollination (1,2 and 3DAP) for *pDD65hp;LIG1GFP*. The box plots cumulate the results obtained for two independent experiments. *LIG1GFP* plants were grown in parallel and quantified as a control. Different symbols represent independent transgenic lines for *pDD35hp;LIG1GFP* (abbreviated *pDD35hp*) and *pDD65hp;LIG1GFP* (abbreviated *pDD65hp*). The following abbreviation are used: central cell (CC), egg cell (EC), synergides (Syn), integuments (Int).

